# Microfluidic Device Facilitates Novel *In Vitro* Modeling of Human Neonatal Necrotizing Enterocolitis-on-a-Chip

**DOI:** 10.1101/2020.11.29.402735

**Authors:** Wyatt E. Lanik, Cliff J. Luke, Lila S. Nolan, Qingqing Gong, Jamie M. Rimer, Sarah E. Gale, Raymond Luc, Shay S. Bidani, Carrie A. Sibbald, Angela N. Lewis, Belgacem Mihi, Pranjal Agrawal, Martin Goree, Marlie Maestas, Elise Hu, David G. Peters, Misty Good

**Author notes:** **Corresponding Author:** Misty Good, MD, MS, Assistant Professor of Pediatrics, Division of Newborn Medicine, Department of Pediatrics, Washington University School of Medicine, St. Louis Children’s Hospital, 660 S. Euclid Ave., Campus Box 8208, St. Louis, MO 63110.

## Abstract

Necrotizing enterocolitis (NEC) is a deadly gastrointestinal disease of premature infants characterized by an exaggerated inflammatory response, dysbiosis of the gut microbiome, decreased epithelial cell proliferation, and gut barrier disruption. Here, we describe a novel *in vitro* model of human neonatal small intestinal epithelium (Neonatal-Intestine-on-a-Chip) that mimics key features of intestinal physiology by utilizing a combination of premature infant intestinal enteroids co-cultured with human intestinal microvascular endothelial cells within a microfluidic device. We used our Neonatal-Intestine-on-a-Chip to recapitulate NEC pathophysiology in an *in vitro* model system of the premature gut inoculated with infant-derived microbiota. This model, also known as NEC-on-a-Chip, emulates the prominent features of NEC, demonstrating significant upregulation of pro-inflammatory cytokines, decreased intestinal epithelial cell markers, reduced epithelial proliferation and disrupted epithelial barrier integrity. NEC-on-a-Chip provides a novel preclinical model of NEC, which may be used as a personalized medicine approach to test new therapeutics for this devastating disease.

## Introduction

Necrotizing enterocolitis (NEC) is a deadly complication of prematurity (1). NEC is characterized by intestinal dysbiosis and an overexaggerated and injurious inflammatory response (1–3). The disease often culminates in decreased epithelial cell proliferation, increased epithelial cell death, a loss of intestinal barrier integrity, and bacterial translocation across the gut barrier (1–3). Despite intensive research with various models of the disease, novel treatment approaches and new preventative strategies for NEC remain lacking. *In vitro* modeling of NEC may accelerate research towards novel preventative strategies and therapeutics; however, current *in vitro* NEC models do not accurately simulate dysbiosis in normal premature intestinal epithelium. For example, rat intestinal epithelial (IEC-6) and human colon adenocarcinoma (Caco-2) cell lines are often used to study molecular pathways and responses to therapeutics in NEC (4–7). However, these monotypic epithelial cell lines do not fully recapitulate the premature intestinal cell architecture and complexity, restricting the physiologic relevance of these models (8).

Other *in vitro* models have used stem cells derived from human small intestinal crypts to produce enteroids (i.e. spheroids or organoids) (9–12). Intestinal organoid models develop all the intestinal epithelial types and have apical-basolateral polarity within a three-dimensional (3D) spherical architecture (9, 13). However, due to their enclosed nature, the ability to study the effect of microbial interactions or therapeutics on the apical epithelial surface is limited. In contrast, a two-dimensional (2D) enteroid monolayer system allows for direct exposure of the apical epithelium to bacteria and therapeutics (14). Despite this advantage, the bacteria in these static systems quickly over-proliferate causing exaggerated epithelial damage and rapid cell death reducing the ability to fully model NEC *in vitro* (15).

Recent advances in microfluidics technology have aided the development of *in vitro* modeling using organ-on-a-chip platforms (15). Intestine-on-a-chip models have been developed to closely simulate the microenvironment of human small intestine through comprehensive cellular differentiation, the formation of 3D villus-like axes, mucus production, continuous luminal flow, and the ability to mimic peristalsis by adding a stretch component (16, 17).

Since dysbiosis is an important feature of NEC, we sought to create a dysbiotic *in vitro* model that recapitulates the main features of the disease. Here, we describe the development of a novel *in vitro* model of a premature infant-derived Neonatal-Intestine-on-a-Chip microphysiological platform to model the interactions between the intestinal epithelium, endothelium, and microbiome. Furthermore, we adapted this Neonatal-Intestine-on-a-Chip model to simulate the pathophysiology of the human disease NEC, called NEC-on-a-Chip, by utilizing a combination of premature infant small intestinal enteroids co-cultured with human intestinal microvascular endothelial cells and patient-derived microbiota, to recreate critical features of premature gut pathophysiology. This novel model system could accelerate the advancement of potential therapeutics for NEC and serve as a tool in precision medicine by the generation of patient-specific treatments.

## Results

### Neonatal-Intestine-on-a-Chip Models Human Small Intestinal Architecture

Due to limitations of current model systems, we sought to develop a Neonatal-Intestine-on-a-Chip model from patient-derived enteroids to better recreate the *in vivo* microenvironment of the newborn small intestine. The developmental progression of intestinal stem cells as enteroids (day 0) to physiologic neonatal intestinal epithelium (day 8) was followed by brightfield microscopy (**Figure 1A**). At day 0, neonatal enteroids appear as discrete organoids. These enteroids were dissociated and seeded into the matrigel-coated microfluidic device (day 1). A confluent monolayer is established by day 3, followed by the appearance of furrows within the epithelium (day 6), and finally advancement to 3D architecture (day 8) with villus-like structures reminiscent of those found *in vivo* (**Figure 1A**).

**Figure 1.**
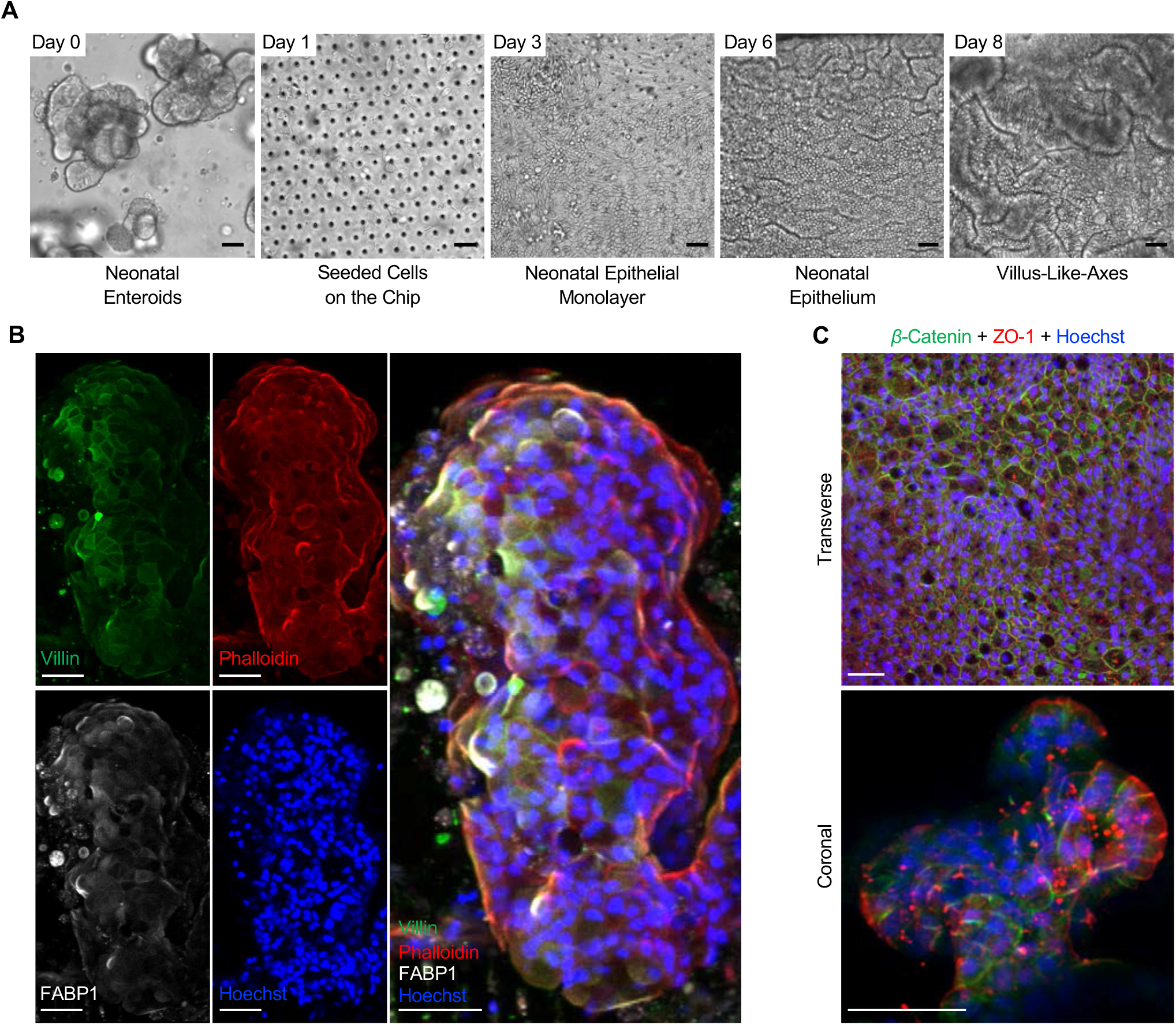
Development of the Neonatal-Intestine-on-a-Chip microfluidic model. Growth progression of Neonatal-Intestine-on-a-Chip is shown by brightfield microscopy in (**A**) beginning with neonatal enteroids at day 0, seeding of stem cells within a microfluidic device on day 1, development of a confluent monolayer by day 3, invaginations of epithelium through day 6, and advancement to villus-like-axes at day 8. Scale bars = 50 μm. (**B**) A representative immunofluorescence image of a deconvoluted coronal cross section of a villus-like formation within the Neonatal-Intestine-on-a-Chip is shown separated (left) and merged (merged) with villin (green, microvilli brush border), phalloidin (red, actin), FABP1 (white, enterocytes), and Hoechst (blue, nuclei). Scale bars = 50 μm. (**C**) Representative immunofluorescent images of Neonatal-Intestine-on-a-Chip epithelium stained with β-catenin (green, basolateral tight junctions), Hoechst (blue), and ZO-1 (red, apical tight junctions) transverse view (top) and coronal cross section (below). Scale bars = 50 μm.

To determine if these villus-like structures have an architecture similar to *in vivo* intestine, the cells within the microfluidic device were immunostained with two molecular markers of intestinal villi: villin, an intestinal brush border marker (green) and fatty acid binding protein 1 (FABP1), an enterocyte marker (white; **Figure 1B**). The villi were counterstained with phalloidin (red) to highlight the actin architecture and Hoescht 33342 to identify nuclei (blue; **Figure 1B)**. Similar to normal human neonatal intestine, these villus-like structures contained an enterocyte brush border with typical actin architecture.

To determine the apical-basolateral polarity within the Neonatal-Intestine-on-a-Chip, cultures were immunostained with primary antibodies directed to the apical tight junction marker, ZO-1 (red), and the basolateral protein, β-catenin (green; **Figure 1C**). Confocal microscopy demonstrated that the ZO-1 marker is concentrated towards the lumen of the Neonatal-Intestine-on-a-Chip, whereas β-catenin is localized towards the semi-permeable membrane (**Figure 1C**). These data suggest that our Neonatal-Intestine-on-a-Chip exhibits 3D cellular architecture with villus-like formations, appropriate cell-cell adhesions, intrinsic cellular polarity, and epithelial cells that mimic human neonatal epithelium.

### Gene Expression Characterization of NEC-on-a-chip

We first evaluated several markers of intestinal gene expression known to be associated with NEC. Analysis of mRNA expression in small intestine tissue from neonates with NEC demonstrated decreased expression of *Leucine-rich repeat-containing G-protein coupled receptor 5* (*LGR5*), a stem cell marker; *Lysozyme,* a Paneth cell marker; and *Mucin 2* (*MUC2*), a goblet cell marker, when compared to healthy control tissue (**Figure 2A**). There was no significant alteration in the expression of the enteroendocrine cell marker, *Chromogranin A* (*CHGA*) in our cohort of patients (**Figure 2A**). Transcriptional analysis of pro-inflammatory cytokines demonstrated that both *interleukin* (*IL*)-*1β* and *IL-8* were upregulated in human NEC tissue compared to controls (**Figure 2B**). Furthermore, we observed a significant reduction in the expression of proliferation markers *Ki67* and *proliferating cell nuclear antigen* (*PCNA*) in NEC tissues when compared to control samples (**Figure 2C**).

**Figure 2.**
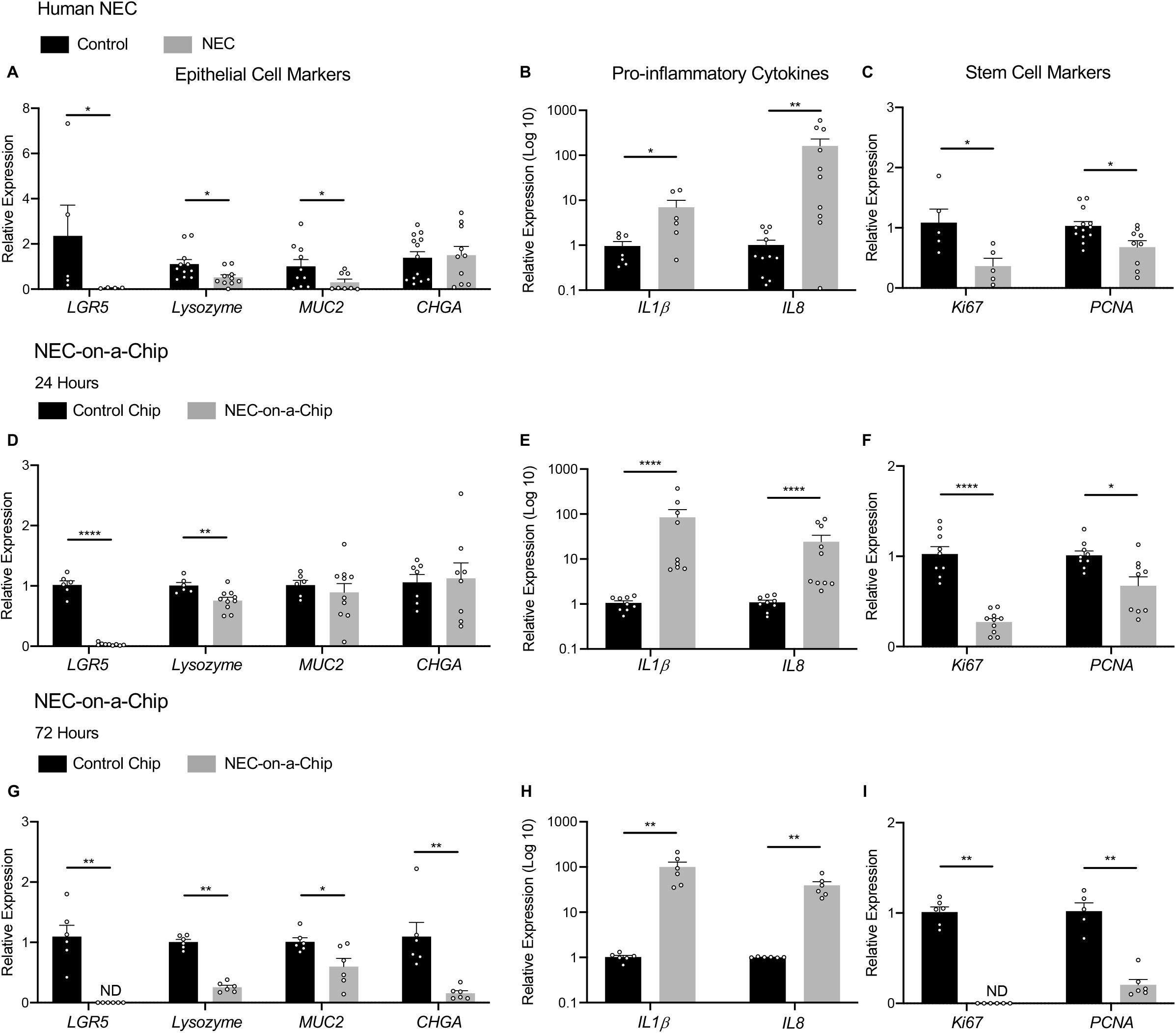
Epithelial cell gene expression is decreased in human NEC and NEC-on-a-Chip. (**A**) Human NEC relative expression of *LGR5* (Control n=5; NEC n=4), *Lysozyme* (Control n=11; NEC n=10), *MUC2* (Control n=10; NEC n=8) and *CHGA* (Control n=13; NEC n=10) epithelial cell markers from neonatal terminal ileal tissue samples with and without NEC were analyzed by qRT-PCR. (**B**) Pro-inflammatory cytokine expression of *IL-1*β (Control n=7; NEC n=6) and *IL-8* (Control n=11; NEC n=10) (**C**) Cell proliferation gene expression of *Ki67* (Control n=4; NEC n=5) and *PCNA* (Control n=13; NEC n=9) was evaluated by qRT-PCR. (**D**) The relative expression of control chips versus NEC-on-a-Chip at 24 hours for *LGR5* (Control n=6; NEC-on-a-Chip n=9), *Lysozyme* (Control n=6; NEC-on-a-Chip n=10), *MUC2* (Control n=6; NEC-on-a-Chip n=10), and *CHGA* (Control n=7; NEC-on-a-Chip n=8). (**E**) Pro-inflammatory cytokines *IL-1*β (Control n=9; NEC-on-a-Chip n=9) and *IL-8* (Control n=9; NEC-on-a-Chip n=10). (**F**) Cell proliferation markers *Ki67* (Control n=9; NEC-on-a-Chip n=10) and *PCNA* (Control n=9; NEC-on-a-Chip n=9). (**G**) The relative expression of control chips versus NEC-on-a-Chip at 72 hours for *LGR5* (Control n=6; NEC-on-a-Chip n=6), *Lysozyme* (Control n=6; NEC-on-a-Chip n=6), *MUC2* (Control n=6; NEC-on-a-Chip n=6), and *CHGA* (Control n=6; NEC-on-a-Chip n=6). (**H**) Pro-inflammatory cytokines *IL-1*β (Control n=6; NEC-on-a-Chip n=6) and *IL-8* (Control n=6; NEC-on-a-Chip n=6). (**I**) Cell proliferation expression at 72 hours by *Ki67* (Control n=6; NEC-on-a-Chip n=6) and *PCNA* (Control n=5; NEC-on-a-Chip n=6) expression were analyzed by qRT-PCR. Data represent mean ± SEM. \**P* < 0.05, \*\**P* < 0.005, \*\*\**P* < 0.0005, \*\*\*\**P* < 0.0001 by Mann-Whitney U test. ND = not detected.

We then utilized our Neonatal-Intestine-on-a-Chip to simulate the *in vivo* intestinal environment during NEC. It is well known that NEC pathogenesis involves intestinal bacterial dysbiosis and an impairment of epithelial barrier integrity which results in bacterial translocation, the induction of pro-inflammatory cytokines, a decrease in epithelial cell proliferation and subsequent cell death, which cannot be corrected by regenerative processes. To model this in our Neonatal-Intestine-on-a-Chip, we co-cultured human intestinal enteroids, endothelial cells, and patient-derived dysbiotic microbiome cultured from a premature infant with severe NEC to create the NEC-on-a-Chip model. In our NEC-on-a-Chip model, we observed significantly downregulated mRNA expression of crypt-associated epithelial cell markers *LGR5* and *Lysozyme* at 24 and 72 hours following introduction of the NEC microbiome when compared to controls (**Figure 2D, G**). At 24 hours, there was no significant difference in the gene expression of intestinal epithelial cell markers *MUC2* and *CHGA* (**Figure 2D**); however, these markers were significantly downregulated at 72 hours (**Figure 2G**).

The NEC bacterial-epithelial co-culture in our NEC-on-a-Chip model resulted in increased expression of pro-inflammatory cytokines *IL-1β* and *IL-8* (**Figure 2E, H**), as well as decreased expression of proliferation markers *Ki67* and *PCNA* (**Figure 2F, I)** at both 24 and 72 hours post-inoculation compared to control chips. Taken together, this data demonstrates the similarities between epithelial cell and inflammatory gene expression in NEC-on-a-Chip and human NEC.

### NEC-on-a-Chip Recapitulates Several Epithelial Features Seen in Human NEC

Confocal immunofluorescence microscopy was used to compare changes in epithelial cell populations seen in human NEC with the NEC-on-a-Chip model 24 hours after introduction of the NEC bacteria. In both human NEC and the NEC-on-a-Chip model, there was a decrease in the number of goblet (Muc2; **Figure 3B, J**), Paneth (Lyz; **Figure 3D, L**), and enteroendocrine (Chromogranin A, CHGA, **Figure 3F, N)** cells when compared to their respective controls of healthy neonatal intestinal tissue (**Figure 3A, C, E, G**) and the Neonatal-Intestine-on-a-Chip (**Figure 3I, K, M, O**), respectively. Confocal microscopy also demonstrated reduction of a proliferation marker, Ki67, in both human NEC tissue samples and NEC-on-a-Chip (**Figure 3H, P)** compared to their respective controls (**Figure 3G, O**). This demonstrates that the epithelial populations decreased in human NEC are also decreased in our NEC-on-a-Chip model.

**Figure 3.**
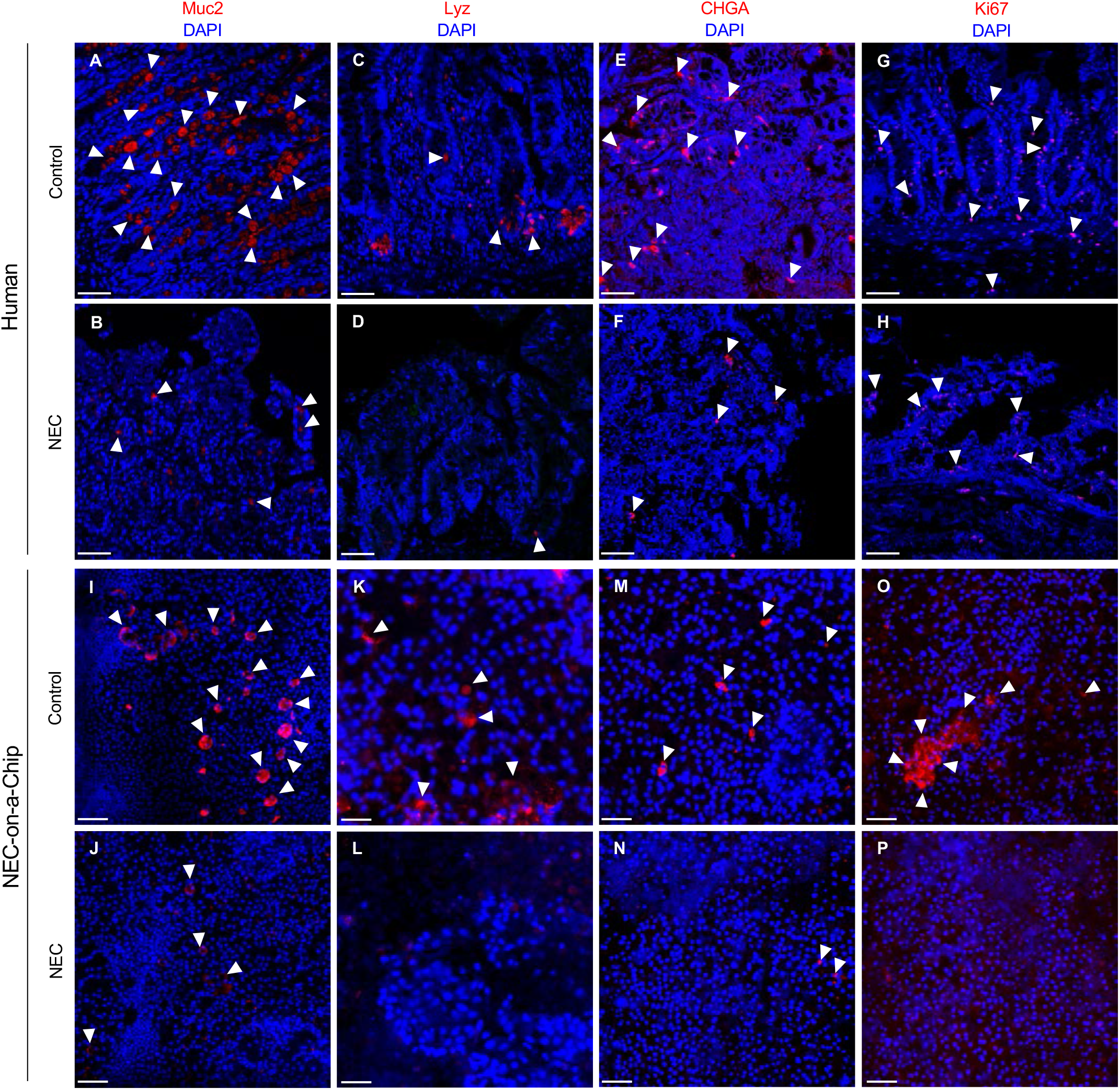
NEC-on-a-Chip recapitulates several features of human NEC. Immunostaining of human control versus NEC ileum (first two rows) and Neonatal-Intestine-on-a-Chip (control) versus NEC-on-a-Chip (last two rows) for the goblet cell marker, Muc2 (red, **A-B, I-J**), Paneth cell marker Lysozyme (Lyz, red, **C-D, K-L)**, Chromogranin A (CHGA, red, **E-F, M-N**) and the proliferation marker Ki67 (red, **G-H, O-P**). White arrows denote representative areas of positive staining. Scale bars = 50 μm.

We next investigated the effects of microbial dysbiosis on the maintenance of epithelial tight junctions in our NEC-on-a-Chip model. Immunofluorescent staining for the epithelial tight junction marker, ZO-1, shows decreased expression at 24-, 48-, and 72-hour timepoints when compared to controls (**Figure 4A, B**). To characterize the degree of epithelial barrier dysfunction in the NEC-on-a-Chip model, we calculated the apparent paracellular permeability (P_app_), a measure of *in vitro* epithelial barrier permeability. The NEC-on-a-Chip demonstrated increasing permeability over 72 hours following NEC bacterial exposure, while the control chips showed no change (**Figure 4C**). This data suggests that we can model the progressive gut barrier dysfunction seen in human NEC in our NEC-on-a-Chip model.

**Figure 4.**
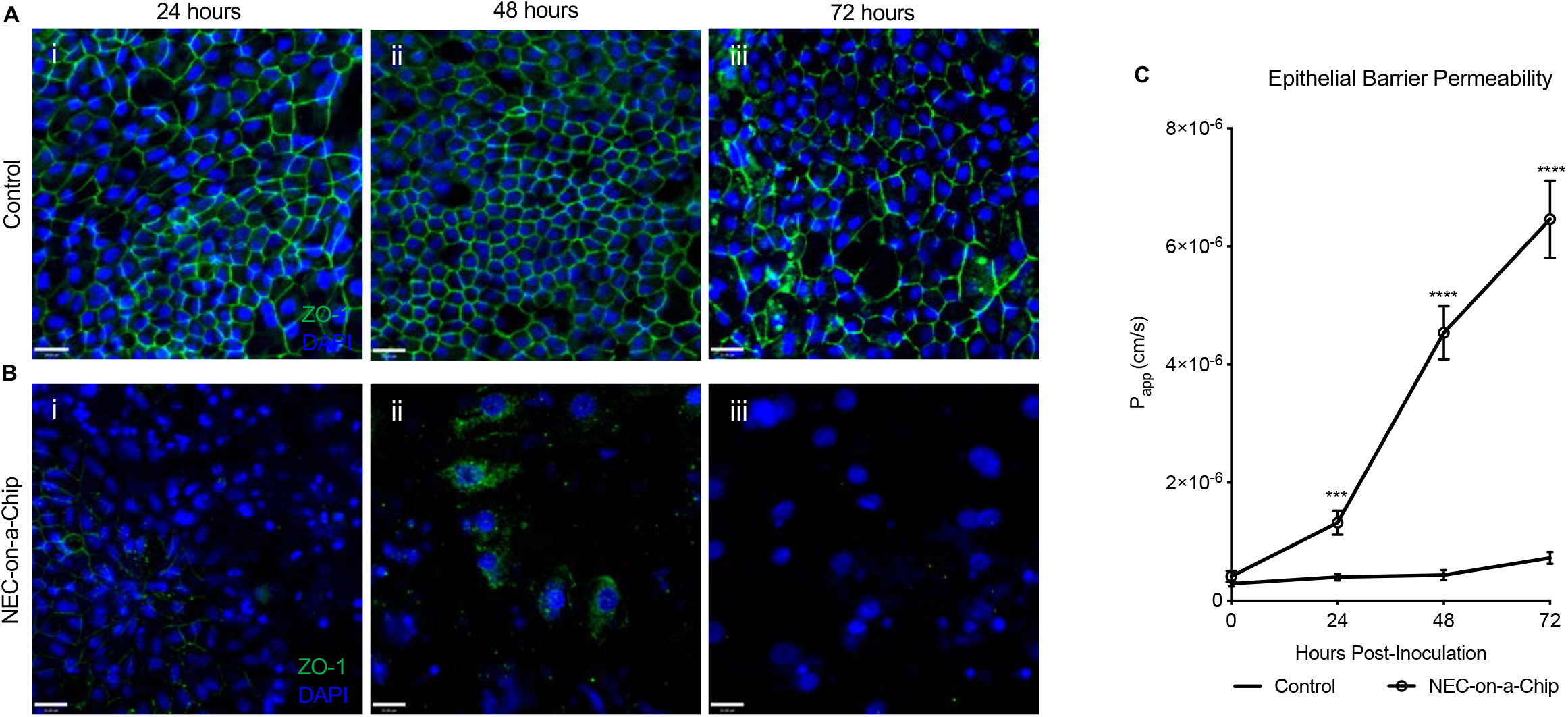
NEC-on-a-Chip has an impaired epithelial barrier. Immunofluorescent staining of tight junction marker ZO-1 in (**A**) control chips and (**B**) NEC-on-a-Chip. (**C**) Apparent paracellular permeability (P_app_) was assessed at 0 (n=23), 24 hours (Control n=12; NEC-on-a-Chip n=12), 48 hours (Control n=6; NEC-on-a-Chip n=6) and 72 hours (Control n=6; NEC-on-a-Chip n=6). Data represent mean ± SEM. \**P* < 0.05, \*\**P* < 0.005, \*\*\**P*<0.0005, \*\*\*\**P* < 0.0001 by unpaired T test. Scale bars = 50 μm.

### Gene Expression Profiling in NEC-on-a-Chip Mimics Key Pathways Induced in Human NEC

To investigate the gene expression profile for NEC-on-a-Chip, we performed RNA-sequencing (RNA-seq) analysis on the small intestine epithelial cells in the NEC-on-a-Chip compared with control chips. A comparison of NEC-on-a-Chip to the controls at 24 hours identified a total of 3354 significantly upregulated and downregulated protein-coding genes. An additional comparison at 72 hours revealed a total of 9515 significantly expressed protein-coding genes. **Figures 5A-B** identifies the most significant pathways modulated in the NEC-on-a-Chip after 24 and 72 hours, respectively. In correlation with the observed increased pro-inflammatory cytokine mRNA expression shown in **Figure 2D** **and** **G**, the RNA-seq data demonstrated upregulated pathways related to inflammation, including the IL-17 signaling pathway, TNF signaling pathway, and cytokine-cytokine-receptor interactions at 24 and 72 hours (**Figure 5A-B**). The heat map in **Figure 5C** displays the 33 most significantly upregulated genes related to pro-inflammatory pathways in the 24-hour NEC-on-a-Chip compared with the control chips, which includes both chemokines, as well as the cytokines IL-1β and IL-6. **Figure 5D** reveals the 34 most significantly upregulated genes related to pro-inflammatory pathways in the 72-hour NEC-on-a-Chips when compared with the control chips.

**Figure 5.**
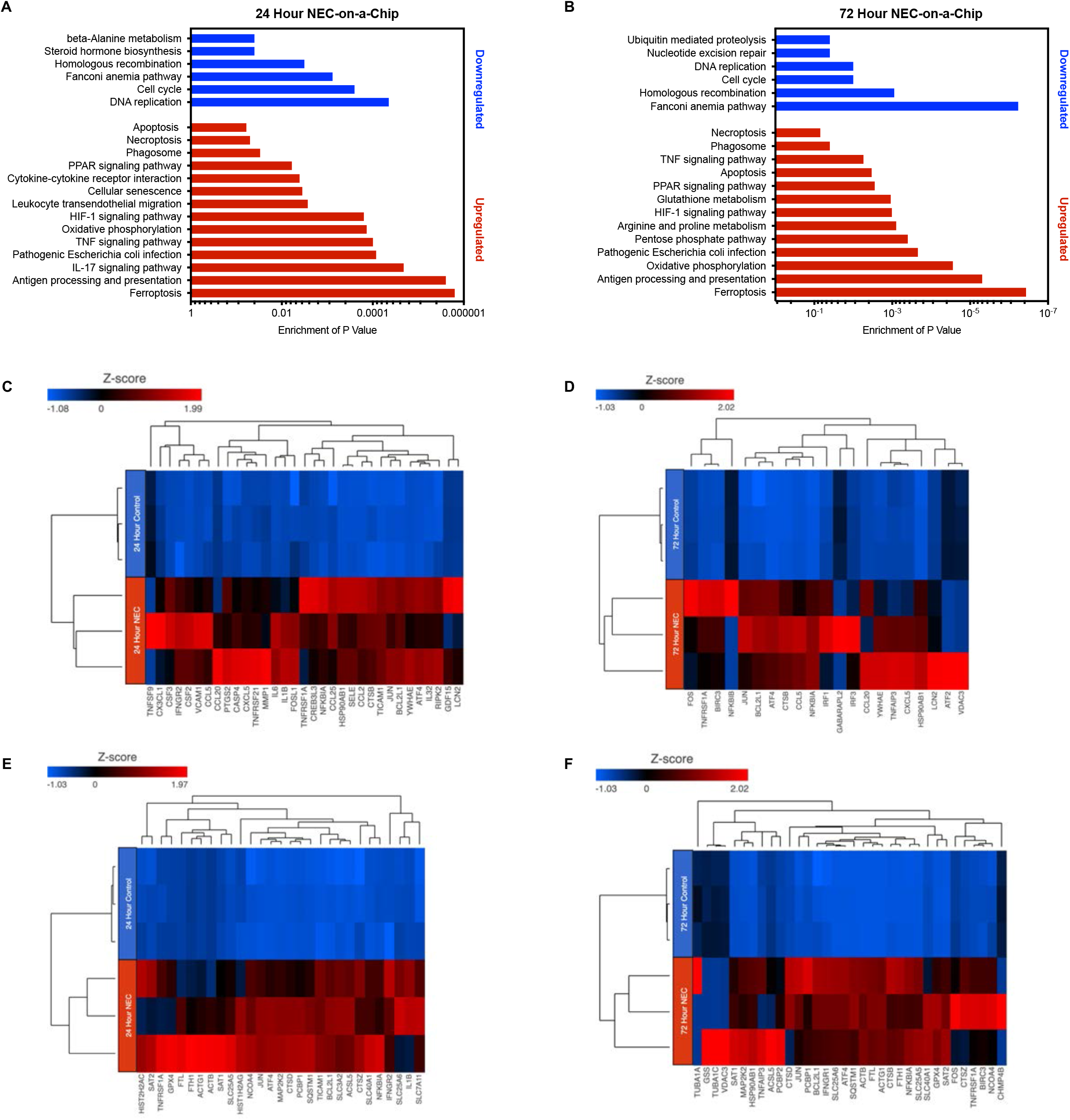
NEC-on-a-Chip contains a distinct profile of gene expression. Functional enriched pathways provide a comparative analysis of NEC-on-a-Chip compared to controls at **(A)** 24 and **(B)** 72 hours. Heat map representation of targeted genes in NEC-on-a-Chip reveals the most significantly upregulated genes in pro-inflammatory pathways at **(C)** 24 and **(D)** 72 hours and in pathways related to cellular death at **(E)** 24 and **(F)** 72 hours.

Importantly, the RNA-seq analysis also revealed significant upregulation of genes involved in cellular death pathways (**Figure 5E-F**). At 24 hours, we observed a significant upregulation of the genes found in the ferroptosis pathway (P <0.0001) in the NEC-on-a-Chip (**Figure 5E**), whereas at 72 hours, significant upregulation in the genes involved in the ferroptosis (P<0.0001) and apoptosis (P<0.005) cell death pathways were observed in the NEC-on-a-Chip when compared to controls (**Figure 5F**). The heat map in **Figure 5E** displays the 29 most significant genes in the apoptosis, ferroptosis, and necroptosis pathways in the 24-hour comparison, and the heat map in **Figure 5F** reveals the 34 most significant genes in these cell death pathways in the 72-hour comparison.

## Discussion

The creation of innovative *in vitro* models for studying human neonatal pathophysiology is paramount to discovering the precise mechanisms involved in disease pathogenesis and developing new epithelial targeted therapeutics. In this study, we demonstrate a novel *in vitro* model of neonatal epithelium that we have termed Neonatal-Intestine-on-a-Chip. Similar to previous adult and pediatric Intestine-on-a-Chip models (16, 17), the Neonatal-Intestine-on-a-Chip overcomes many limitations of traditional 2D and 3D intestinal cell culture techniques.

The Neonatal-Intestine-on-a-Chip produces villus-like 3D architecture with robust epithelial cell differentiation into goblet, Paneth, and enteroendocrine cells proportionate to intestine of a premature infant. Epithelial polarity is preserved as evidenced through staining of the apical tight junctions with ZO-1 and basolateral proteins with β-Catenin. The apical epithelial surface is easily accessible for exposure to experimental stimuli, such as bacteria or treatment with therapeutics. Additionally, the dynamic luminal flow with simulated peristaltic-like movements mimics intestinal peristalsis seen in humans. These features make the Neonatal-Intestine-on-a-Chip model an accurate *in vitro* reflection of the premature intestinal microenvironment for the study of neonatal disease.

The Neonatal-Intestine-on-a-Chip was inoculated with enteric bacteria from a neonate with severe NEC to emulate the characteristic intestinal dysbiosis seen in NEC. This novel NEC-on-a-Chip system accurately simulates several key features of the neonatal disease, including exaggerated inflammatory cytokine production, decreased epithelial cell populations and reduced proliferation, as well as progressive permeabilization of the gut barrier.

Similar to the pro-inflammatory signature of human NEC, the NEC-on-a-Chip model demonstrated increased expression of *IL-1β* and *IL-8* at both 24 and 72 hours following inoculation with the NEC bacteria. Additionally, RNA sequencing analysis of the NEC-on-a-Chip epithelium, found increased expression of several cytokines, chemokines, and inflammatory markers including *IL-1β*, *IL-6*, *CXCL5*, and *Lipocalin 2* (*LCN2*). These findings suggest a pro-inflammatory microenvironment in NEC-on-a-Chip that is comparable to human NEC.

The pro-inflammatory environment seen in human NEC is associated with significant epithelial cell death and decreased epithelial cell populations. RNA sequencing analysis showed upregulation of genes involved in the apoptosis, ferroptosis, and necroptosis pathways in the NEC-on-a-Chip compared to controls. The NEC-on-a-Chip model also demonstrated decreased mRNA expression of stem cell marker *LGR5* by 24 hours and complete loss of expression by 72 hours, reflecting the trends seen in human NEC tissue from both this and previously reported studies (18, 19). Paneth cell (lysozyme), goblet cell (MUC2), and chromogranin A (CHGA) populations were decreased in both our NEC tissue and NEC-on-a-Chip. This is consistent with previous NEC literature demonstrating a decrease in these cell populations during the disease (20–22).

The role of enteroendocrine cells in NEC pathogenesis is not well established. Our evaluation of human tissue samples showed no difference in mRNA expression, however there was an apparent decrease in the CHGA immunofluorescent staining between NEC and control samples. Immunofluorescence staining for CHGA in NEC-on-a-Chip at 24 hours revealed an observable difference between NEC-on-a-Chip and controls. Furthermore, the NEC-on-a-Chip model did show decrease in *CHGA* mRNA expression at the 72-hour timepoint. Additional studies will be needed to look for the presence of late enteroendocrine cell injury in NEC and elucidate the physiologic significance of this phenomenon.

Our NEC-on-a-Chip model also accurately reflected the decreased cell proliferation seen in the neonatal disease (4, 5, 18, 19, 23). The mRNA expression of both *Ki67* and *PCNA* proliferation markers were decreased in human NEC tissue and NEC-on-a-Chip samples compared to controls. Immunofluorescence imaging confirmed decreased Ki67 staining of both human NEC tissue and NEC-on-a-Chip samples compared to controls, indicating a decreased capacity for epithelial proliferation.

The loss of tight junctions in NEC contributes to a leaky gut barrier (24, 25) allowing bacterial translocation and increasing the risk of sepsis and death (18, 26, 27). Previous studies have found that *ZO-1* mRNA expression is decreased in NEC tissue (24, 25, 28). We also demonstrated a decrease in the expression of the tight junction ZO-1 as well as its internalization by confocal microscopy and further demonstrated that the apparent paracellular permeability of the NEC-on-a-Chip system increases with time.

The NEC-on-a-Chip model simulates many key *in vivo* features of NEC. It is a practical *in vitro* platform for studying NEC pathophysiology and testing candidate therapeutics. Since this model can be produced with patient-specific epithelium, it could be the first step towards developing a personalized medicine approach to study and treat NEC.

## Methods

### In vitro culture of human neonatal enteroids

De-identified human ileal tissue was collected from neonates undergoing required resection in accordance with Washington University School of Medicine in St. Louis approved guidelines and regulations. Tissue was processed as previously described in VanDussen KL *et al* (29). In brief, neonatal small intestinal tissue was washed with Dulbecco’s Modified Eagle’s Medium/F12 (DMEM/F12, Invitrogen) with 4-(2-hydroxyethyl)-1-piperazineethanesulfonic acid (HEPES, Corning) supplemented with 10% heat inactivated fetal bovine serum (Sigma), 2 mM L-glutamine (Gibco), 100 units mL^−1^ penicillin and 0.1 mg mL^−1^ streptomycin (Sigma) to inactivate endogenous proteases. The tissue was collagenase I digested, mechanically dissociated to release crypts, filtered, washed, and centrifuged for crypt isolation. Crypts were suspended in Matrigel growth factor-reduced matrix (Corning), which was polymerized at 37°C for 20 minutes. Enteroids were cultured in enteroid growth media consisting of a 50% vol/vol mixture of Advanced DMEM/F12 (Invitrogen) and conditioned media from an L cell line producing Wnt, R-spondin and Noggin ligands (L-WRN) (30) supplemented with 10 μM Y-27632 (R&D), 10 μM SB-431542 (R&D), 100 units mL^−1^ penicillin and 0.1 mg mL^−1^ streptomycin. Enteroid growth media was replaced every 2 to 3 days. Enteroids were passaged every 5 to 7 days by incubating in 0.25% trypsin 0.5 mM phosphate buffered saline-ethylenediaminetetraacetic acid (PBS-EDTA, Gibco) solution for 3 minutes at 37°C, followed by mechanical dissociation. Enteroids between passage number 7 and 20 were used to seed the chips.

### Neonatal-Intestine-on-a-Chip culture

Two chambered microfluidic chips were obtained from Emulate Inc. (Boston, MA) and treated as previously described (16) with minor modifications. Briefly, chips were activated following the manufacturer’s instructions using ER1 and ER2 solutions (Emulate Inc.). The top chamber of the chips was coated with 200 μg ml^−1^ type IV collagen (Sigma) and 100 μg ml^−1^ Matrigel growth factor-reduced matrix in PBS (Gibco), while the bottom chamber was coated with 200 μg ml^−1^ type IV collagen and 30 μg ml^−1^ fibronectin (Sigma) in PBS. The chips were then placed in a humidified incubator at 37°C overnight. Cultured neonatal human enteroids were dissociated and resulting enteroid fragments were resuspended at a density of 6×10^6^ cells mL^−1^ in chip expansion media (50% vol/vol mixture of Advanced DMEM/F12 and L-WRN with 10 μM Y-27632, 10 μM SB-431542, 10 nM human [Leu15]-gastrin I (Sigma), 500 nM A83-01 (Sigma), 10 mM HEPES, 100 units mL^−1^ penicillin, 0.1 mg mL^−1^ streptomycin, 1x B-27 Supplement (Gibco), 1x N-2 Supplement (Gibco), 1 mM N-acetylcysteine (Sigma), 50 ng mL^−1^ murine epidermal growth factor (EGF, PeproTech) and 10 mM Nicotinamide (Sigma)). The top channel of the chip was loaded with 30 uL of resuspended cells (~180,000 cells per chip) and maintained at 37°C overnight. The following day seeded chips were washed with pre-warmed expansion media to remove unattached cells. Continuous perfusion with expansion media was initiated through the top and bottom channels with a flow rate of 30 μL hr^−1^. Once the neonatal epithelium reached confluency, repeated peristaltic-like membrane contractions were initiated at 10% strain and 0.2 Hz.

Upon appearance of villus-like axes (day 6), Human Small Intestinal Microvascular Endothelial Cells (HIMECs; Cell Biologics) were resuspended at a density of 6×10^6^ cells mL^−1^ in Microvascular Endothelial Cell Growth Medium-2 media (EGM-2MV, Lonza). The bottom channel of the chip was washed with EGM-2MV and loaded with 6 μL of HIMEC cell suspension (~36,000 cells per chip). The chips were inverted for 2 hours at 37°C to facilitate attachment of the HIMECs to the semi-permeable membrane, opposite the confluent neonatal epithelium. After adhesion of HIMECs to the membrane, continuous flow at 30 μL hr^−1^ was reinitiated with expansion media and EGM-2MV media to the epithelial and endothelial channels, respectively.

### Bacterial culture

An aliquot of an enteric bacterial slurry isolated from a patient with severe, surgical NEC was obtained as previously described (31). This bacterial slurry was grown from 10 μL of a frozen glycerol stock incubated in 2 mL Luria-Bertani (LB) broth (Sigma) at 37°C. The bacterial slurry was diluted in chip expansion media to 7×10^8^ CFU mL^−1^.

### NEC-on-a-Chip model

Antibiotic-containing cell culture media was removed from the media reservoirs 24 hours prior to bacterial inoculation and media reservoirs were subsequently rinsed with sterile PBS. Antibiotic-free expansion media and antibiotic-free EGM-2MV media were added to the top and bottom channel reservoirs, respectively. Where indicated, 30 μL of the bacterial slurry (7×10^8^ CFU mL^−1^) was added after 24 hours to the top channel of the chip (2.1×10^7^ CFU per chip). The chips remained under static condition for 30 minutes to facilitate bacterial adherence to the apical side of the neonatal epithelium. The top channel was flushed with antibiotic-free expansion media to remove unattached bacteria and the chips were continuously perfused with antibiotic-free media at 30 μL hr^−1^. Chips were flushed at an increased rate of 1000 μL hr^−1^ for 6 minutes every 24 hours to remove unattached bacteria.

### Intestinal permeability assessment

To measure apparent paracellular permeability (P_app_), 50 μg mL^−1^ Cascade Blue (3 kDa, Thermo Fisher) was added to the antibiotic-free expansion media in the top epithelial channel with a flow rate of 30 μL hr^−1^. Every 24 hours, the effluents from the epithelial and endothelial channels were collected. The concentration of Cascade Blue that diffused across the membrane into the endothelial channel effluent was quantified with a Varioskan LUX multimode microplate reader (Thermo Fisher) with fluorescence intensities of 400 nm and 425 nm. The following formula was used to calculate P_app_ (cm s^−1^) as in (16):

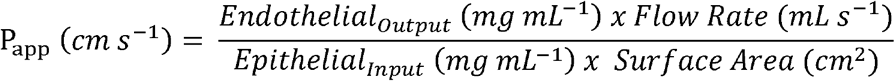

where *Endothelial*_*Output*_ is the concentration of Cascade Blue from the endothelial channel effluent; *Flow Rate* is the fluid flow rate of the media through both channels; *Epithelial*_*Input*_ is the concentration of Cascade Blue originally added to the epithelial channel media; and *Surface Area* is the area of the channel where co-culture occurs (cm^2^).

### RNA-Sequencing and Analysis

Bulk RNA-sequencing was performed on small intestine epithelial cells obtained from the NEC-on-a-Chip and control chips by the Genome Technology Access Center (GTAC) at the Washington University School of Medicine in St. Louis. Total RNA was isolated using TRIzol™. RNA samples with a RIN of at least 7 were used to generate libraries by GTAC using the RNA zero kit (Roche). Samples were prepared according to library kit manufacturer’s protocol, indexed, pooled, and sequenced on an Illumina HiSeq. Basecalls and demultiplexing were performed with Illumina’s bcl2fastq software and a custom python demultiplexing program with a maximum of one mismatch in the indexing read.

To generate heat maps and for the pathway analysis, mRNA expression was analyzed using Partek^®^ Flow^®^ software. Adapters were removed and reads were aligned to the MM10 assembly with STAR 2.7.3a and quantified to Ensembl transcript release 91 with an average coverage of 10x. Reads were normalized to total counts per million. Features containing fewer than 10 reads were not included in the analysis. Sequencing performance was assessed for the total number of aligned reads, the total number of uniquely aligned reads, and features detected. The ribosomal fraction, known junction saturation, and read distribution over known gene models were quantified. Ribosomal genes and genes not expressed in the smallest group size minus one sample greater than one count-per-million were excluded from further analysis. The performance of all genes was assessed with plots of the residual standard deviation of every gene to their average log-count with a robustly fitted trend line of the residuals. Differential expression analysis was then performed to analyze for differences between groups and the results were filtered for only those genes with p-values less than or equal to 0.05 and a fold change over 2.0 was applied between any pairwise comparisons.

### RNA isolation, reverse transcription, and qRT-PCR

Epithelial RNA was isolated from the top channel of the chips by administration of TRIzol™. cDNA was synthesized using QuantiTect Reverse Transcription Kit (Qiagen) per the manufacturer’s instructions. Quantitative real-time PCR (qRT-PCR) was performed using SsoAdvanced™ Universal SYBR^®^ Green Supermix (Bio-Rad) in conjunction with a CFX Connect™ Real-Time PCR Detection System (Bio-Rad). Primers were used as shown in **Table 1** and relative expression levels were normalized to *RPLO* as a housekeeping gene.

**Table 1.**
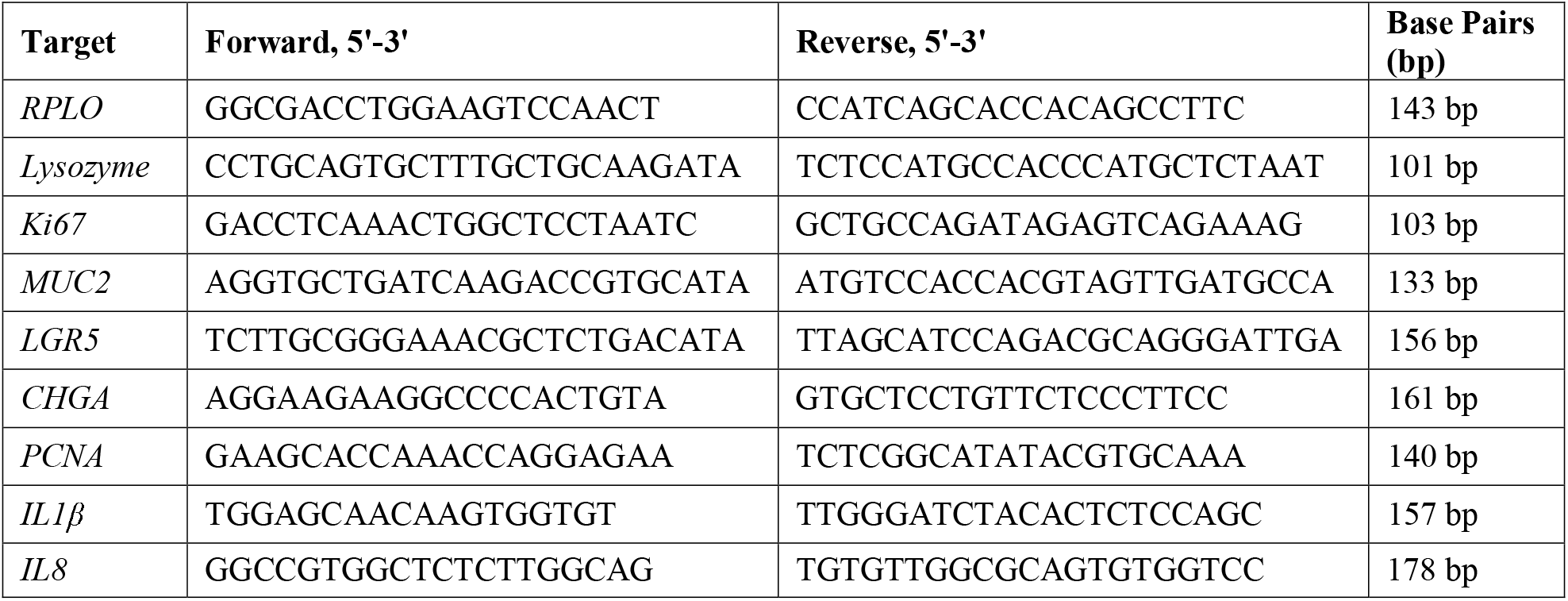
List of qRT-PCR primers.

### Immunofluorescence

Immunofluorescence staining was performed on fixed, paraffin-embedded neonatal ileal tissue. Briefly, neonatal tissue slides were deparaffinized in xylene, rehydrated in isopropanol and boiled in antigen unmasking solution (Vector Laboratories). Slides were then blocked with 1% bovine serum albumin (BSA, VWR) and 10% normal donkey serum (NDS, Sigma) in 1x Tris Buffered Saline (Boston BioProducts) with 0.1% Tween-20 (Sigma) (TBST). Primary antibodies (**Table 2**) were diluted 1:100 in blocking solution and secondary antibodies (Life Technologies) were diluted 1:200 in blocking solution. Nuclear staining was performed using Hoechst 33342 (Invitrogen) and slides were mounted with Prolong Gold (Thermo Fisher). Slides were imaged on either a Zeiss Axio Observer.Z1 using an EC Plan-Neofluar 10x objective (Zeiss) with either a 1x tube-lens or 1.6x tube-lens optical variable adapter (Zeiss) or a white light laser Leica SP8X tandem scanning confocal microscope (Leica Microsystems) with a 40×1.3 NA oil objective (Leica Mircrosystems). Images were processed using either ZEN pro, blue edition (v2.3, Zeiss) or LASX (Leica Microsystems) software, respectively.

**Table 2.** List of primary antibodies used for immunofluorescence.

| **Antibodies** | **Isotype** | **Manufacturer** | **Catalog #** | **Dilution** |
| --- | --- | --- | --- | --- |
| Lysozyme | goat | Santa Cruz | sc-27958 | 1:100 |
| Ki67 | goat | Santa Cruz | sc-7846 | 1:100 |
| MUC2 | rabbit | Santa Cruz | sc-15334 | 1:100 |
| CHGA | rabbit | Abcam | ab15160 | 1:100 |
| Villin | mouse | Santa Cruz | sc-58897 | 1:100 |
| ZO-1 | rabbit | Zymed | 40-2200 | 1:100 |
| PCNA | mouse | Santa Cruz | sc-56 | 1:100 |
| β-Catenin | mouse | Santa Cruz | sc-7963 | 1:100 |
| FABP1 | rabbit | Novus Biologicals | NBP1-87695 | 1:100 |
| Phalloidin | N/A | Invitrogen | R415 | 1:100 |

Immunofluorescent staining of the chips was performed as previously described (16). In brief, chips were fixed in 4% paraformaldehyde (Alfa Aesar), permeabilized with 0.1% Triton X-100 (Sigma) and blocked in 10% NDS in PBS. Chips were stained with primary antibodies (**Table 2**) diluted in 5% NDS in PBS, and secondary antibodies diluted 1:200 in 5% NDS in PBS, and either Hoechst or DAPI. Where indicated, coronal cross sections of the chips were hand-cut with a razor blade with approximately 300 μm thickness. Coronal cross sections were then imaged in an imaging dish with PBS.

A white light laser Leica SP8X tandem scanning confocal microscope was used to image the chips with a 25x water objective (Leica Microsystems). Images were acquired every 0.5 μm in the *z*-plane to render 3D images. Images were generated and analyzed using LASX and Volocity (v6.3.5; Quroum Technologies) software. Where indicated, 3D deconvolution was performed using Huygens Essential, Version 19.10 (Scientific Volume Imaging).

### Light Microscopy

The epithelial structure of the top channel of the chip was assessed over time with brightfield microscopy using a Zeiss Axio Observer.Z1 in combination with either an EC Plan-Neofluar 10x objective or a Plan-Apochromat 10x objective (Zeiss) and either a 1x tube-lens or a 1.6x tube-lens optical variable adaptor (Zeiss). Live-cell, representative *z*-plane images were obtained at 24-hour time intervals with temperature and carbon dioxide control settings at 37°C and 5% CO_2_, respectively. Images were processed using ZEN 2.3 pro, blue edition software.

### Statistics

Statistical analyses with Mann-Whitney U test and two-tailed unpaired T tests were performed where indicated using GraphPad Prism software version 8.0. Outliers were identified using the ROUT method. Statistical significance was accepted at a p value of <0.05.

### Study approval

Deidentified premature infant small intestine was obtained from infants undergoing surgical resection for NEC or for non-inflammatory conditions (controls). All specimens were processed as discarded tissue with a waiver of consent approval of the Washington University in St. Louis School of Medicine Institutional Review Board (IRB Protocol numbers 201706182 and 201804040).

## Author Contributions

Conceptualization, W.E.L., C.J.L., S.E.G. and M.G.; Methodology, all authors; Investigation, all authors; Writing – Original Draft, W.E.L., and M.G.; Writing – Review & Editing, all authors; Funding Acquisition, C.J.L., L.S.N and M.G.; Resources, M.G.; Supervision, M.G.

## Acknowledgements

This manuscript was supported R01DK118568 (MG), R01DK114047 (CJL) and 5T32HD043010 (LSN) from the National Institutes of Health, the St. Louis Children’s Hospital Foundation (MG), the Children’s Discovery Institute of Washington University and St. Louis Children’s Hospital (CJL, MG), and the Department of Pediatrics at Washington University School of Medicine, St. Louis.

## Notes

**Funding:** This manuscript was supported R01DK118568 (MG), R01DK114047 (CJL) and 5T32HD043010 (LSN) from the National Institutes of Health, the St. Louis Children’s Hospital Foundation (MG), the Children’s Discovery Institute of Washington University and St. Louis Children’s Hospital (CJL, MG), and the Department of Pediatrics at Washington University School of Medicine, St. Louis.

**Conflicts of interest:** MG has received sponsored research agreement funding from Astarte Medical Partners and Takeda Pharmaceuticals. None of these sources had any role in this study.

### Competing Interest Statement

MG has received sponsored research agreement funding from Astarte Medical Partners and Takeda Pharmaceuticals. None of these sources had any role in this study.

## References

1. Neu J, Walker WA. Necrotizing enterocolitis. The New England journal of medicine 2011;364(3):255–64.

2. Morrow AL et al. Early microbial and metabolomic signatures predict later onset of necrotizing enterocolitis in preterm infants. Microbiome 2013;1(1):13.

3. Nino DF Sodhi CP, Hackam DJ. Necrotizing enterocolitis: mechanisms and management. Nature Reviews Gastroenterology and Hepatology 2016;In press.

4. Good M et al. Breast milk protects against the development of necrotizing enterocolitis through inhibition of Toll-like receptor 4 in the intestinal epithelium via activation of the epidermal growth factor receptor. Mucosal Immunol 2015;8(5):1166–79.

5. Good M et al. Amniotic fluid inhibits Toll-like receptor 4 signaling in the fetal and neonatal intestinal epithelium. Proceedings of the National Academy of Sciences 2012;109(28):11330–11335.

6. Leaphart CL et al. A critical role for TLR4 in the pathogenesis of necrotizing enterocolitis by modulating intestinal injury and repair. Journal of immunology 2007;179(7):4808–20.

7. Wang C et al. Human Milk Oligosaccharides Activate Epidermal Growth Factor Receptor and Protect Against Hypoxia-Induced Injuries in the Mouse Intestinal Epithelium and Caco2 Cells. J. Nutr. 2020;150(4):756–762.

8. Sun H, Chow EC, Liu S, Du Y, Pang KS. The Caco-2 cell monolayer: usefulness and limitations. Expert Opin Drug Metab Toxicol 2008;4(4):395–411.

9. Sato T et al. Single Lgr5 stem cells build crypt-villus structures in vitro without a mesenchymal niche. Nature 2009;459(7244):262–5.

10. Zachos NC et al. Human Enteroids/Colonoids and Intestinal Organoids Functionally Recapitulate Normal Intestinal Physiology and Pathophysiology. J Biol Chem 2016;291(8):3759–66.

11. Afrazi A et al. Toll Like Receptor 4-mediated Endoplasmic Reticulum Stress in Intestinal Crypts Induces Necrotizing Enterocolitis [Internet]. The Journal of biological chemistry [published online ahead of print: February 11, 2014]; doi:10.1074/jbc.M113.526517

12. Werts AD et al. A Novel Role for Necroptosis in the Pathogenesis of Necrotizing Enterocolitis. Cell Mol Gastroenterol Hepatol 2020;9(3):403–423.

13. Stelzner M et al. A nomenclature for intestinal in vitro cultures. Am. J. Physiol. Gastrointest. Liver Physiol. 2012;302(12):G1359–1363.

14. Senger S et al. Human Fetal-Derived Enterospheres Provide Insights on Intestinal Development and a Novel Model to Study Necrotizing Enterocolitis (NEC). Cell Mol Gastroenterol Hepatol 2018;5(4):549–568.

15. Williamson IA et al. A High-Throughput Organoid Microinjection Platform to Study Gastrointestinal Microbiota and Luminal Physiology. Cell Mol Gastroenterol Hepatol 2018;6(3):301–319.

16. Kasendra M et al. Development of a primary human Small Intestine-on-a-Chip using biopsy-derived organoids. Sci Rep 2018;8(1):2871.

17. Kasendra M et al. Duodenum Intestine-Chip for preclinical drug assessment in a human relevant model. Elife 2020;9. doi:10.7554/eLife.50135

18. Neal MD et al. Toll-like receptor 4 is expressed on intestinal stem cells and regulates their proliferation and apoptosis via the p53 up-regulated modulator of apoptosis. The Journal of biological chemistry 2012;287(44):37296–308.

19. Li B et al. Impaired Wnt/β-catenin pathway leads to dysfunction of intestinal regeneration during necrotizing enterocolitis. Cell Death Dis 2019;10(10):743.

20. Coutinho HB et al. Absence of lysozyme (muramidase) in the intestinal Paneth cells of newborn infants with necrotising enterocolitis. J. Clin. Pathol. 1998;51(7):512–514.

21. Sodhi CP et al. Intestinal epithelial Toll-like receptor 4 regulates goblet cell development and is required for necrotizing enterocolitis in mice. Gastroenterology 2012;143(3):708-18 e1–5.

22. McElroy SJ et al. Tumor necrosis factor receptor 1-dependent depletion of mucus in immature small intestine: a potential role in neonatal necrotizing enterocolitis. American journal of physiology. Gastrointestinal and liver physiology 2011;301(4):G656–66.

23. Sodhi CP et al. Toll-like receptor-4 inhibits enterocyte proliferation via impaired beta-catenin signaling in necrotizing enterocolitis. Gastroenterology 2010;138(1):185–96.

24. Moore SA et al. Intestinal barrier dysfunction in human necrotizing enterocolitis. J. Pediatr. Surg. 2016;51(12):1907–1913.

25. Liu D et al. Mucins and Tight Junctions are Severely Altered in Necrotizing Enterocolitis Neonates. Am J Perinatol [published online ahead of print: May 23, 2020]; doi:10.1055/s-0040-1710558

26. Hackam DJ, Good M, Sodhi CP. Mechanisms of gut barrier failure in the pathogenesis of necrotizing enterocolitis: Toll-like receptors throw the switch. Seminars in pediatric surgery 2013;22(2):76–82.

27. Ford HR. Mechanism of nitric oxide-mediated intestinal barrier failure: insight into the pathogenesis of necrotizing enterocolitis. Journal of pediatric surgery 2006;41(2):294–9.

28. Bein A, Eventov-Friedman S, Arbell D, Schwartz B. Intestinal tight junctions are severely altered in NEC preterm neonates. Pediatr Neonatol 2018;59(5):464–473.

29. VanDussen KL et al. Development of an enhanced human gastrointestinal epithelial culture system to facilitate patient-based assays. Gut 2015;64(6):911–920.

30. Miyoshi H, Ajima R, Luo CT, Yamaguchi TP, Stappenbeck TS. Wnt5a potentiates TGF-β signaling to promote colonic crypt regeneration after tissue injury. Science 2012;338(6103):108–113.

31. Good M et al. Lactobacillus rhamnosus HN001 decreases the severity of necrotizing enterocolitis in neonatal mice and preterm piglets: evidence in mice for a role of TLR9. American journal of physiology. Gastrointestinal and liver physiology 2014;306(11):G1021–32.

